# A tale of two vineyards: parsing site-specific differences in bacterial and fungal communities of wine grapes from proximal vineyards, and their changes during processing in a single winery

**DOI:** 10.1101/2025.03.05.641676

**Authors:** Reid G. Griggs, Lena Flörl, Michael Swadener, Rodrigo Hernández-Velázquez, David A. Mills, Nicholas A. Bokulich

## Abstract

Wine is a microbial product, naturally transformed through fermentation by a consortium of fungi and bacteria that originate from the vineyard and the cellar, in addition to any microorganisms that are intentionally inoculated. Previous work has shown that grapevine-associated microbiota follow distinct biogeographic patterns, associated with climate and soil properties, and that even neighboring vineyards can harbor distinct microbial communities, but it is unclear whether these differences persist when controlling for variations in farming practices, cultivar, and climate, and whether site-specific microbial profiles change during processing in the winery. Here we investigated the bacterial and fungal microbiota of fruit pre- and post-harvest from two nearby vineyards planted to a single variety, geographically close to one another, and farmed the same way, and then processed in a single winery. These communities subtly changed during processing yet retained distinct site-specific signatures, indicating a partial contribution of the winery environment to the microbiota of grape must and juice pre-fermentation. We also profiled the microbiota of key microbial sources in the winery environment, including fruit flies and processing equipment, demonstrating that the microbiota at these sites reflect contact with plant material, harbor communities distinct from fruit, and appear to partially contribute to the fermentation assemblage, especially via the contribution of fermentative yeasts that are rare or missing in the vineyard environment. These results bolster previous reports of site-specific microbial signatures in winegrowing and make a first estimation of the changes to the grape-associated microbiome during early processing.

**Importance:** Native wine fermentations are driven by microbes carried over from the vineyard or introduced in the winery. In this study, we tracked the microbiome dynamics of wine fermentations from two Chardonnay vineyards planted in close proximity to examine the relative contribution of vineyard- and winery-resident microbiota on microbial succession during wine fermentation. By tracking microbial changes from vineyard to winery, we show that the winery environment, including processing equipment and fruit flies, contributes to the fermentation microbiome but does not override vineyard-specific microbial differences. These findings support the concept of *microbial terroir* and highlight the importance of vineyard microbiomes in shaping wine fermentation. This work advances our understanding of how microbial diversity influences wine production and provides insight into the ecological dynamics of fermentation. By identifying key microbial sources and their contributions, this study lays the groundwork for future microbiome research in viticulture and wine making.

## Introduction

Microorganisms are integrally involved in wine production, with the microbial inhabitants of vineyards and wineries being entirely responsible for fermentation of grape juice in traditional “native” wine fermentations. Microbial metabolism plays a critical role in shaping the resulting wine quality, and hence the observation that grapevine-associated microbiota exhibit biogeographic patterns associated with climate, soil characteristics, and other factors [1], [2], [3], [4], [5] raises the exciting possibility that regional variation in grape-associated microbiota contribute to regional and site-specific variations in wine quality, a hypothesis referred to as *microbial terroir* [6]. This follows conceptually from the idea of *terroir*, that wines are influenced by the regional and site-specific growing conditions for grapevines in general.

Numerous studies have reported spatiotemporal variation in grape and wine microbiota in virtually every major winegrowing region. The same grapevine cultivar can harbor different microbial communities when the fruit is grown in geographically distinct vineyards [1], [7], [8], [9], [10], [11], as well as phenotypically distinct strains of *Saccharomyces cerevisiae* [8], [12], and the grape microbiome pre-fermentation predictively correlates with the differential production of metabolites post-fermentation [2], [11], [13], [14]. This highlights the potential role of autochthonous microorganisms in regional wine production, but many questions still remain regarding which biotic and abiotic factors shape the site-specific microbial communities associated with grapevines, and how resilient these communities are during processing in the winery.

The winery environment has also been shown to be a medium for the transfer of microorganisms, and although a resident population of fermentative yeasts can be detected in the cellar, a bolus of microbial biomass enters the winery during harvest [15]. A large percentage of previously undetected taxa from vineyard surveys has been reported to appear in musts using culture-independent methods [3]. However, it is unclear from prior work whether these microorganisms originated from the winery environment, and to what degree the grape microbiota fluctuate during fruit processing.

While ample evidence points to the structuring of microbial communities associated with fruit and must by region and variety [5], [16], [17], the strength of the link between these two parts of the winegrowing continuum, and the relative influence of the winery are not clear. Establishing an understanding of how microbial communities change through fruit processing in a production winery is conceptually critical to understanding the extent of the previously reported features of *microbial terroir*. Here, we investigated, for the first time, how the grape microbiota fluctuate during processing and what the potential contribution of winery-resident microbiota may be, including microbes from fruit flies. Fruit from two different vineyards was sampled pre-harvest and then tracked through processing, and the winery environment was surveyed as well. A primary motivation for this study was to assess whether vineyards planted to the same variety, farmed in the same manner, in close proximity to one another with no major topographic features between them harbored distinct microbial communities, and whether these microbial signatures persisted through grape processing to contribute site-specific microbial assemblages to the fermentation.

## Methods

### Vineyard sites

Microbial communities were sampled from fruit in two teaching vineyards in Davis California named RMI (38.53155, −121.7542) and Tyree (38.52675, −121.78946) on 8/7/2013. The vineyards are located 3.15 km apart, and both planted to Chardonnay, cordon trained with vertical shoot positioning, and farmed conventionally. RMI vineyard is directly adjacent to the UC Davis teaching winery and Interstate 80, while Tyree vineyard is set between a creek and the UC Davis Airport, surrounded by farmland. In addition to Chardonnay, Tyree is planted with mono-varietal blocks of Colombard, Grenache Blanc, Grenache Noir, and Cabernet Sauvignon, among other varieties. These five varieties were sampled in this study to examine variation within the vineyard, and to follow changes in composition during processing. All varieties are cordon trained with vertical shoot positioning and farmed conventionally. Clone, rootstock, soil, and spacing information for each variety sampled is shown in Table 1. Tyree Chardonnay was planted to a mixture of loamy alluvial land and the Reiff soil series, classified as coarse-loamy, mixed non-acid thermic Mollic Xerofluvents. RMI and the other varieties in Tyree are planted to the Yolo soil series, described as fine-silty, mixed, superactive, thermic Fluventic Haploxerepts.

**Table 1.** Vineyard site information.

| Vineyard | Variety | Clones | Rootstocks | Soil | Row Spacing | N Samples* |
| --- | --- | --- | --- | --- | --- | --- |
| Tyree | Chardonnay | 4, 6, 29 | 101-14, 5BB | Reiff, Loamy alluvial | 8x12 (ft) | 36 |
|  | Colombard | 2 | 101-14 | Yolo | 8x12 (ft) | 36 |
|  | Grenache Blanc | 141 | 101-14 | Yolo | 8x12 (ft) | 48 |
|  | Cabernet Sauvignon | 8 | Unknown | Yolo | 8x12 (ft) | 37 |
|  | Grenache Noir | 224 | 3309C | Yolo | 8x12 (ft) | 25 |
| RMI | Chardonnay | 4, 45, 66 | 420A | Yolo | 7x11(ft) | 60 |

\* N samples = The number of unique vines sampled in this study.

### Sample Collection

Berries were aseptically removed from vines at regular intervals to cover the entire block of each vineyard and placed in sterile Whirl-Pak sampling bags (Weber Scientific, Hamilton, NJ, USA). Specifically, 8 vines each from 3 rows were sampled from Tyree vineyard, and 12 vines each from 5 rows were sampled from RMI vineyard. Samples were taken on different dates for each variety so that they were collected within two weeks prior to harvest, so that samples would be collected at approximately similar ripeness; Chardonnay was sampled on 8/7/13 in both vineyards, Cabernet Sauvignon on 9/5/13 and 9/25/13, Colombard and Grenache Blanc were sampled on 9/5/13 and 9/16/13, and Grenache Noir was sampled on 9/30/13. Samples were collected from equidistant vines on either side of a vineyard row, while walking down the row to best represent the entire space of the vineyard. Five samples of crushed fruit (must) were also aseptically collected in sterile Whirl-Pak sampling bags (Weber Scientific, Hamilton, NJ, USA) from the destemmer during the processing of each lot. Must samples were placed immediately on ice and were handled like the berry samples, juice was pressed off and samples were frozen at - 80 °C until processing. Samples were also directly collected from the press for each lot, at regular 10-minute intervals reflecting changes in the must/juice microbiota across the press cycle, and frozen immediately at −80 °C.

Prior to processing, on 8/15/13 crush equipment was aseptically swabbed with sterile cotton-tipped swabs (Puritan Medical, Guiliford, ME) as previously described by Bokulich et. al 2013 [15]. Briefly, swabs were moistened with PBS and streaked across an approximately 10 cm square area in overlapping S-strokes in an attempt to cover the entire surface and use the entire surface of the swab. Swab heads were aseptically snapped off into 1.5 mL polyethylene tubes and immediately placed on ice. Samples were then stored in a −80 °C freezer until processing. Two samples were taken from each of the following pieces of crush equipment: the hopper, elevator belt, destemmer, pump, press bladder, press pan rail, and press pan.

Flies were collected from McPhail traps [18] placed in five different locations of the fermentation hall at the UC Davis Teaching Winery during the harvest of 2013. Traps were sterilized with 10 % by volume bleach solution between uses, and baited before each use with pressed pomace from a Cabernet Sauvignon fermentation that had been inoculated with a commercial strain of *S. cerevisiae*. To separate flies from bait, circular filters were cut, autoclaved, and aseptically fitted to traps after baiting. Traps were collected within 6-24 hours of being set, on dates ranging from 9/27/2013 to 10/24/2013. Upon collection, traps were stored at - 20 °C until processing to kill flies. Up to 25 flies were randomly selected from each trap and collected aseptically, placed in sterile DNA free 1.5mL microcentrifuge tubes and immediately stored at −80 °C until extraction. Whole flies were pooled and processed by trap as is common [19].

A total of 242 grapes samples were collected from RMI and Tyree vineyards as detailed in Table 1; 60 must or juice samples at different stages of processing; 68 fruit fly samples; and 13 equipment swabs collected from different fruit-contacting surfaces of the destemmer, crusher, and press.

### Sample Processing

Grape samples were aseptically crushed immediately upon return to the laboratory, and juice was frozen at −80°C until processing. Grapes from each vineyard were manually harvested (Chardonnay: 8/15/13; Cabernet Sauvignon: 9/25/13; Colombard: 9/26/13; Grenache Blanc: 9/27/13; Grenache Noir: 10/4/13), and each block was processed individually at the Robert Mondavi Institute for Food and Wine Science Winery (University of California, Davis). Fruit processing included dumping each of 5 bins into a hopper, where fruit came into contact with the elevator belt, feeding into the destemmer. Fruit pumped with a must pump into a bladder press and pressed with 60-70-minute ramping cycles.

Samples from fruit and musts/juices were processed as described previously [1], and winery equipment swabs were processed as previously described [20]. Briefly, must samples were thawed and centrifuged at 4,000 x *g* for 15 min, washed in ice-cold PBS, and suspended in 200 uL DNeasy lysis buffer, with an added 40 mg/mL lysozyme, and then incubated at 37 °C for 30 min. Samples from winery swabs were thawed and aseptically transferred to ZR-96 Fecal DNA MiniPrep Kit (Zymo Research, Irvine CA) tubes. All samples were then processed according to the manufacturer’s protocol, with the addition of a 2 min maximum speed bead beater cell lysis step using a FastPrep-24 bead beater (MP Bio, Solon, OH).

### Microbiome marker-gene amplicon sequencing and data analysis

Amplification and sequencing were completed as previously described [2]. Specifically, the V4 domain of the 16S rRNA gene was amplified using the universal primer pair F515/R806 [21]. The internal transcribed spacer 1 (ITS1) loci were amplified with the BITS/ B58S3 primer pair [22]. Extracted DNA was checked for quality with gel electrophoresis and a NanoDrop spectrophotometer (Thermo Fischer Scientific, DE, USA). Amplicons of the V4 and ITS regions were separately pooled in equimolar ratios, purified using the Qiaquick spin kit (Qiagen), and submitted to the UC Davis Genome Center DNA Technologies Core Facility (Davis, Ca, USA) for Illumina paired-end library preparation, and 250-bp paired-end sequencing on an Illumina MiSeq (Illumina, Inc. CA, USA).

Marker-gene amplicon sequence data were analyzed using the QIIME 2 microbiome bioinformatics framework version 2024.10 [23]. Raw Illumina fastq files were demultiplexed using the QIIME 2 plugin for cutadapt [24] and denoised using the plugin for DADA2 [25] to generate ASVs (amplicon sequence variants). Bacterial V4 reads were truncated at 140 nts (forward) and 200 nt (reverse) based on quality scores; primers were removed by trimming 21 nt (forward reads) and 23 nt (reverse reads). ASVs were classified taxonomically using the q2-feature-classifier plugin with a weighted naive Bayes taxonomic classifier [26], [27] using the SILVA reference database version 138.1 nr99 [28]. For fungal ITS data, only forward reads were used due to low-quality reverse reads. ITS reads were truncated at the first position with a Q-score < 10, and ASVs otherwise processed as described for bacterial V4 reads. Fungal taxonomic classification we performed using q2-feature-classifier against the Unite ITS database (version 10.0; fungi and eukaryotes; 99% OTUs) [29]. ASVs were filtered to remove sequences not assigned at the class level, ASVs observed in only one sample or fewer than 100 times, and any ASVs that were assigned to kingdoms Viridiplantae, Metazoa, or Stramenopila (for ITS sequences) or to Chloroplast, Mitochondria, or Eukaryota (for V4 sequences). A total of 23 negative controls (blanks) were included in both the V4 and ITS sequencing runs; all had no or very few reads and were removed prior to downstream analysis.

Diversity analyses, including calculations of alpha-diversity (richness and Shannon entropy) and beta-diversity (Jaccard distance and Bray-Curtis dissimilarity) were made using the q2-diversity and q2-kmerizer [30] plugins in QIIME 2. Beta diversity estimates were performed based on ASV composition, as well as the constituent k-mers in these ASVs, using a k-mer decomposition technique prior to diversity analysis [30]. This enables diversity estimates that are based on sequence similarity between communities, so that distances are effectively weighted by the degree of shared sequence space; genetically distinct communities will have larger distances, and genetically similar communities will be more similar even if they are composed of distinct strains of the same species. Feature tables were normalized by sampling at even depths (without replacement) prior to diversity estimates and statistical testing: 1000 reads per sample for fungal ITS reads and 500 reads per sample for bacterial reads. Principal coordinates analysis (PCoA) was used to reduce dimensionality prior to visualization. Visualizations were generated using seaborn [31]. Statistical testing for differences in alpha and beta diversity were performed using ANOVA (as implemented in q2-longitudinal [32] and statsmodels [33] and PERMANOVA adonis tests [34] (as implemented in q2-diversity and r-vegan [35]), respectively.

Random forests supervised classification models [36] implemented in scikit-learn [37] were trained to predict sample type, vineyard, and variety based on bacterial and fungal community composition. Models were trained using 500 trees and 10-fold nested cross-validation as implemented in the q2-sample-classifier plugin [26]. Evenly sampled, total-sum-scaled ASV abundances were used as training data.

### Network analysis

To reconstruct the ASV co-occurrence network, we first merged the ASV count data from the V4 and ITS datasets, resulting in a matrix of 281 samples. This matrix was used to generate an artifact for the QIIME2 plugin q2-makarsa [38]. The network was inferred using the Flashweave algorithm [39] within q2-makarsa, employing the sensitive mode with a Q-value threshold of <0.01. To refine the network, only positive edges were retained, and edges with a weight < 0.3 were removed.

A bipartite network was constructed to represent the relationships between samples and ASVs. To ensure biological relevance, this network was filtered to include only ASV nodes present in the co-occurrence network, as these ASVs exhibit significant associations and potential biological interactions. For visualization, ASVs were grouped by species, with edge weights representing the number of ASVs connecting different samples. Additionally, samples were categorized into four groups: Berry, Must, Fruit Flies, and Equipment Swabs. A Gephi visualization [40] was generated while maintaining equal node weights for species and samples. To optimize the graphical representation and highlight closely associated sample categories, the Yifan Hu force-directed algorithm was applied [41].

## Results

We sought to investigate microbial community compositions and transmission routes from the vineyard into the winery, to examine whether site-specific microbial communities persist through fruit processing up until the point of initiation of fermentation. Samples were collected from grapes in the vineyard prior to harvest; of grapes during processing and after crushing and pressing (must and juice; referred to only as must throughout for simplicity); from winery equipment surfaces; and from fruit flies captured within the winery (Figure 1). After quality filtering, 7,048,852 ITS reads in 281 samples representing 1,310 unique amplicon sequence variants (ASVs) and 471 taxa remained, and 2,851,978 V4 reads representing 881 unique ASVs and 451 taxa across 383 samples were observed. We traced the journey of the fruit from the vineyard through the winery to test variances in community composition (i) between vineyards pre-harvest; (ii) between grape varieties pre-harvest; and (iii) between sample types pre- and post-harvest to examine community similarity between grapes, must, and winery sources to (iv) examine source-sink relationships.

**Figure 1.**
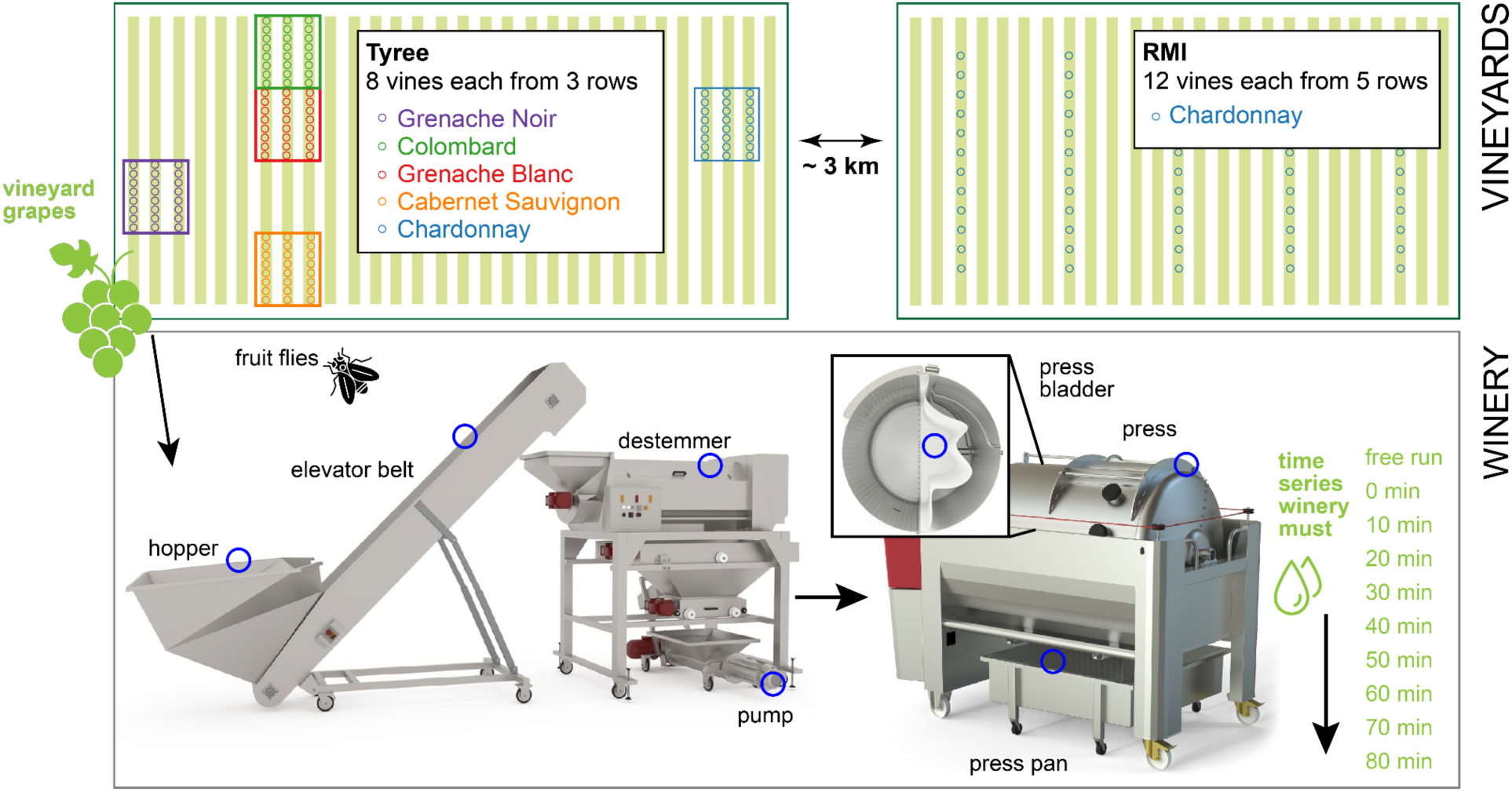
Overview of the sampling strategy. Grape samples were collected from two vineyards, Tyree and RMI, both of which were planted with Chardonnay. Additionally, samples from other grape varieties were collected from the Tyree vineyard (Grenache Noir, Colombard, Grenache Blanc, Cabernet Sauvignon). All grapes were processed in the same winery, where swabs were taken from various pieces of equipment throughout the processing workflow, including the hopper, elevator belt, destemmer, pump, press bladder, press pan rail, and press pan (indicated by blue circles). Fruit flies were also collected as potential vectors of microbial communities. Finally, samples of the resulting grape must were obtained.

### Grape microbiota exhibit vineyard site and varietal variation

First, we compared bacterial and fungal communities of grape samples collected at commercial ripeness from two nearby vineyards in Davis, CA, as well as pressed must samples from the winery where grapes from both vineyards were processed. The vineyards were farmed under identical conditions to confirm previous findings that grape microbiota exhibit significant differences in diversity and microbial community composition based on (i) site and (ii) variety. We again observed significant differences in both bacterial and fungal beta diversity (PERMANOVA P-values = 0.001) (Table 2, Figure 2, Figure S1) and alpha diversity (ANOVA P-values < 0.001) between grapevine cultivars; this varietal effect was observed in both grapes collected in the vineyards and musts from the winery (Figure S2). Significant differences were observed with the unweighted metric Jaccard distance, which measures distance between communities based on the presence/absence of different features, and with Bray-Curtis dissimilarity, which measures community dissimilarity based on the presence and abundance of different features. Multi-way PERMANOVA tests comparing vineyard sites, varieties, and berry vs. must samples indicate that grape variety is the strongest explanatory factor (Table 2), with particularly strong associations with bacterial community composition (Jaccard PERMANOVA R2 = 0.191, P = 0.001). Vineyard and sample type are significant factors, but associated with a much smaller degree of variance in the bacterial and fungal communities (Table 2).

**Figure 2.**
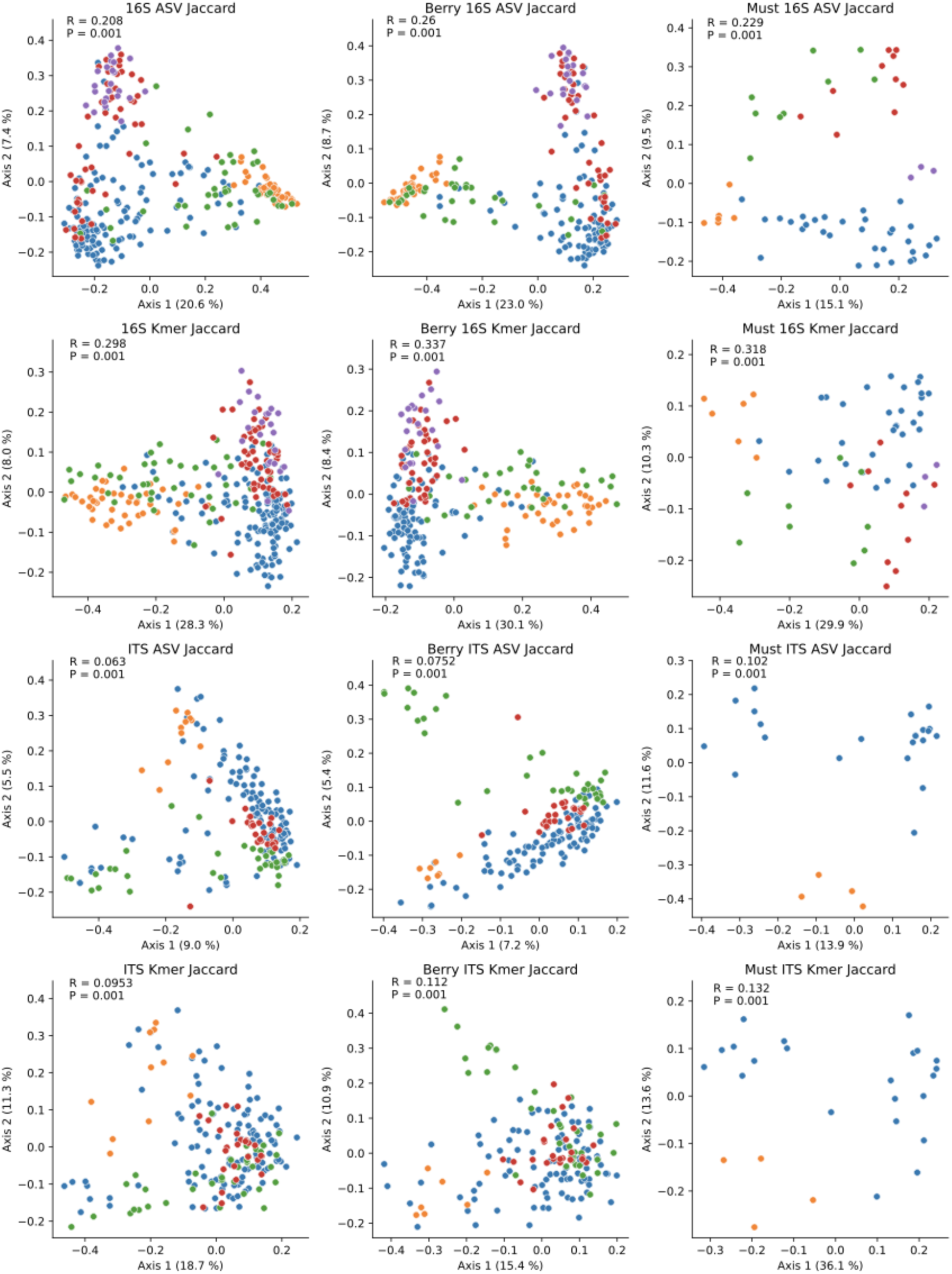
Grape varieties retain distinct microbiota composition through processing. Jaccard distance PCoA Plots of grape and must (left), grape berry (middle) and must samples (right) based on bacterial (16S) and fungal (ITS) composition; and colored by variety. (Blue = Chardonnay; Orange = Cabernet Sauvignon; Green = Colombard; Red = Grenache Blanc; Purple = Grenache Noir. Inset: one-way PERMANOVA R2- and P-values.)

**Table 2.**
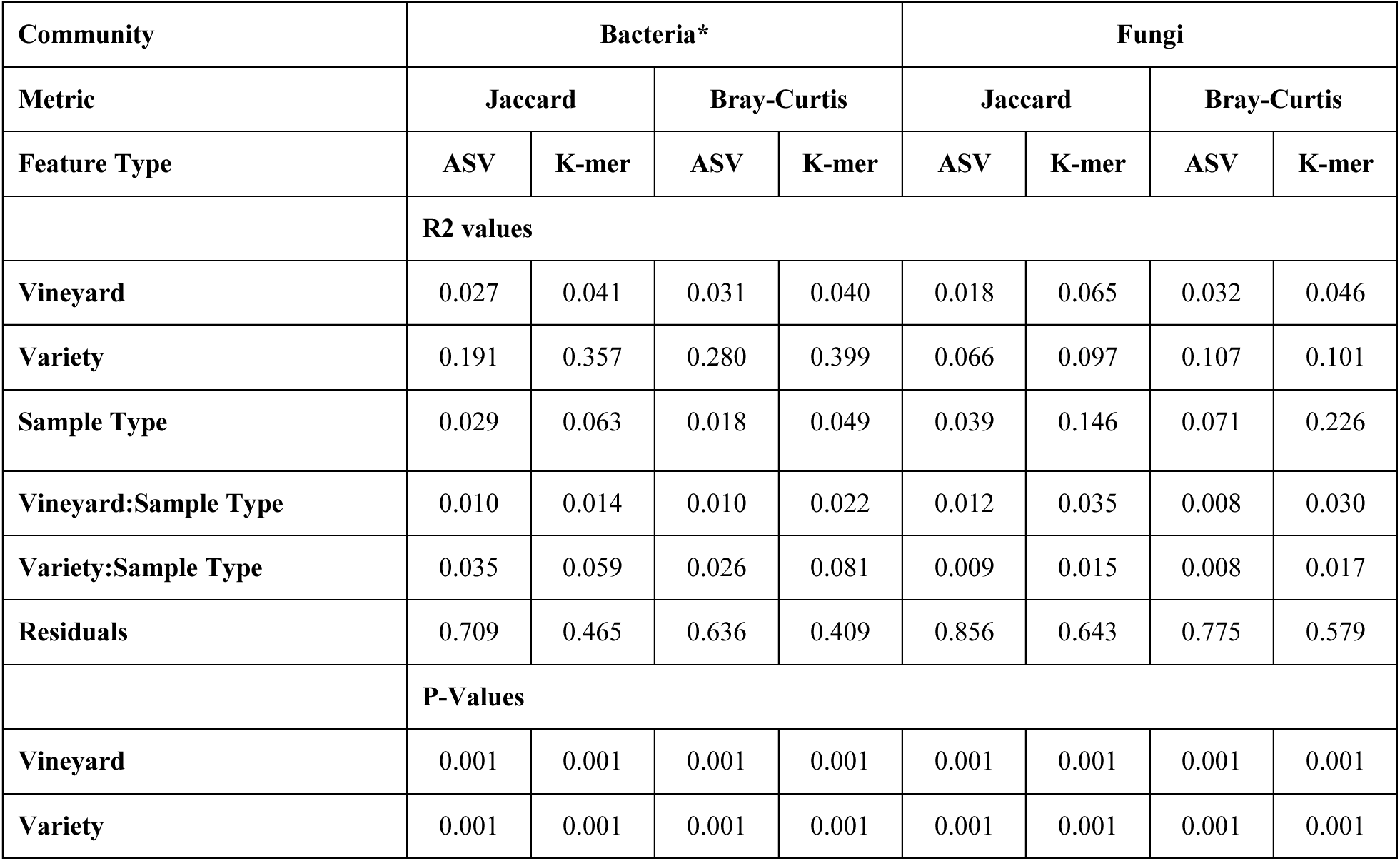
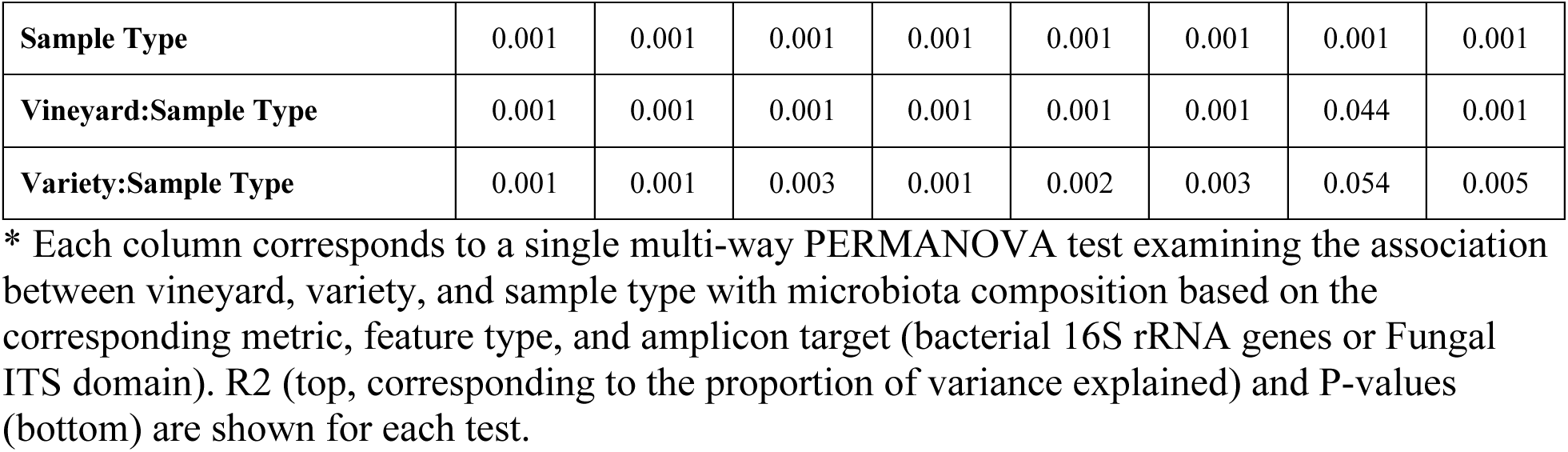
PERMANOVA tests for associating bacterial and fungal community composition with vineyard, grape variety, and sample type (grape berries or must).

The degree of variation explained by microbiota composition can be determined from the PERMANOVA R2-value, which indicates the amount of variation explained by a given factor. This difference is most significant when comparing only grape samples (highest PERMANOVA R-value = 0.574, Figure S1), but even when combining grape and must samples together, strong variety effects are observed (Figure 2, Figure S1; leftmost columns), indicating that these variety effects are strongly preserved through grape processing and pressing, and indeed persist into the must (Figure 2, Figure S1, rightmost columns). Thus, fungal and bacterial community signatures present in the vineyard are delivered to the fermentation tanks more or less intact, and any fermentative yeasts and bacteria present in the vineyard may be expected to contribute to fermentation, particularly in uninoculated fermentations.

The strongest R2-values were observed with beta diversities based on k-mer composition (Figure 2, Figure S1), which provides a measure of community similarity based on shared genetic signatures [30]. This suggests that differences between grape varieties are driven by differences in genetically diverse species of bacteria and fungi. Additionally, large effect sizes are seen for bacterial community comparisons (as high as R=0.574, Figure S1), indicating that bacterial community composition is distinct between grape varieties; fungal communities show lower but still significant and strong effect sizes (as high as R=0.203, Figure S1), suggesting that fungal community composition varies strongly between grape varieties, but to a more moderate degree. To determine which ASVs were differentially abundant between grapevine varieties, we trained Random Forest supervised classification models (via nested 10-fold cross-validation) to predict the variety based on bacterial and fungal composition. These models demonstrated high accuracy for predicting grapevine variety (0.81 ± 0.07 % accuracy; micro-AUC 0.98; Figure S3), corroborating the strongly significant differences in beta diversity. The top 20 important features identified by the models (indicating relative strength for predicting which variety a sample belongs to) are primarily filamentous fungi that are common residents in vineyards, including several species of *Alternaria*, *Aureobasidium pullulans*, *Cladosporium cladosporioides*, and several bacteria with no known fermentative ability (Figure S4).

We also assessed whether site-specific differences in bacterial and fungal diversity of grapes were present in these vineyards, reflecting previous findings of differences between vineyards at small spatial scales [2], [42]. Given the apparent effects of grapevine variety, we compared a single variety — Chardonnay — grown in the Tyree and RMI vineyards. Results indicate that beta diversity is significantly different between these vineyards (PERMANOVA P = 0.001; Figure 3), though effect sizes are somewhat lower than observed for varietal differences (minimum R = 0.0367 for 16S ASV Jaccard distance; maximum = 0.272 for ITS k-mer Bray-Curtis dissimilarity; Figure 3). Effect sizes are larger for fungal communities and for k-mer-based metrics, suggesting that genetically diverse fungi play a more significant role in shaping site-specific community compositions in vineyards that are grown under similar management regimes and very similar climate conditions, to a single variety (∼3 km apart in a topographically homogenous region of central California). Bacterial and fungal ASV richness and Shannon diversity were also significantly higher in Tyree than in RMI (Figure S5).

**Figure 3.**
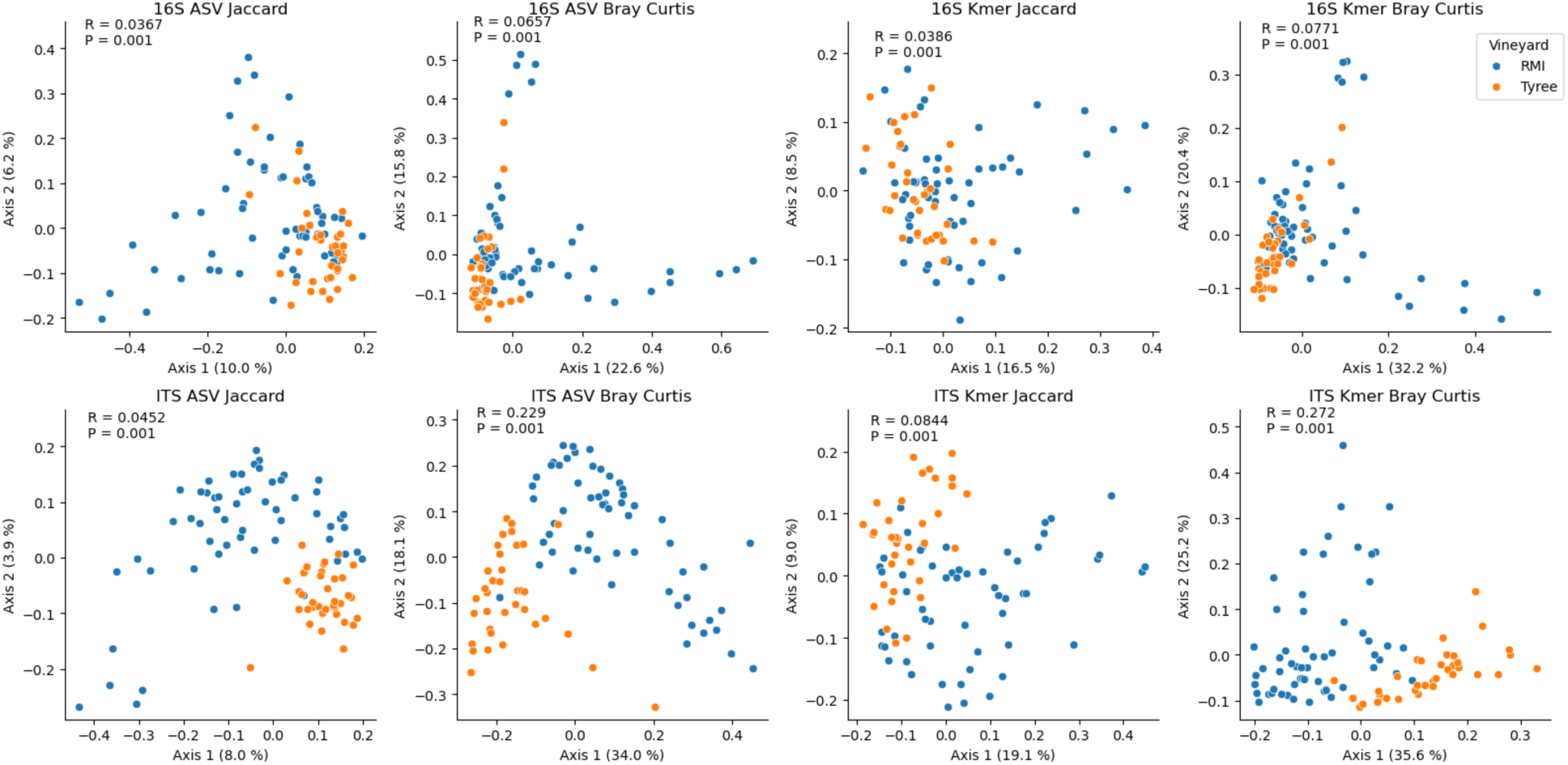
Chardonnay grapes from proximal vineyards harbor significantly different microbial communities. Jaccard distance PCoA Plots of Chardonnay samples collected from RMI (blue) and Tyree (orange) vineyards, based on bacterial (16S) and fungal (ITS) composition. Inset: One-way PERMANOVA R2- and P-values.

Random Forest classifiers indicated very high predictive accuracy for differentiating these vineyards (accuracy = 0.935 ± 0.052; micro-AUC = 0.98; Figure S6). The top 20 most important ASVs again included mostly filamentous fungi and non-fermentative bacteria (Figure S7). However, many ASVs assigned to fermentative yeast species could be detected within the grape samples; 20 different unique *Hanseniaspora* ASVs were detected (belonging to *H. uvarum* and *H. vineae*), and many were frequently detected in Tyree but this species was nearly absent in RMI (Figure S8).

### Must microbiota reveal imprint of winery microbiota but retain site-specific signatures

Next, we examined how the bacterial and fungal communities of grapes pre-harvest differ from those of grape musts following harvest and processing. PERMANOVA tests indicate significant differences between grape and must microbiota when controlling for vineyard site and grape variety (all varieties tested) (Table 2). In general, the amount of variance explained by sample type is much lower than the amount explained by grape variety and comparable to the amount explained by vineyard site; the one exception is for fungal k-mer-based beta diversity, which registers high variance explained for both Jaccard distance (R2=0.146, P=0.001) and Bray-Curtis dissimilarity (R2=0.226, P=0.001). High beta diversity with these metrics and not ASV-based metrics implies an emergence of genetically diverse fungi in the must, leading to the presence of unique k-mers that drive these observed differences. It should be noted that the multi-way PERMANOVA tests included all grape varieties, and so site-specific effects are conflated with varietal effects in this test (as only Chardonnay was collected from the RMI vineyard), so the tests described in the previous section, which compared only Chardonnay samples from the two vineyards, should be referred to for examining site-specific effects. Next, we directly compared the microbiota of grapes, must, and winery sources (equipment and fruit flies) to assess the degree of similarity between vineyard and winery ecosystems. These sites all strongly demonstrate significant differences in beta diversity (Figure 4) and alpha diversity (Figure 5). Although these differences are mostly driven by large differences between fruit fly microbiota and all other sample types, subtle significant differences are observed between grape and must samples, including higher bacterial ASV richness (mean 7.69, FDR-corrected P_ANOVA_ < 0.001) and higher fungal ASV Shannon H (Mean 0.398, FDR-corrected P_ANOVA_ = 0.009) in must than in grapes (other comparisons were insignificant). This indicates that bacterial alpha diversity slightly increases during fruit processing, suggesting possible influx of microorganisms from winery sources, and fungal evenness decreases (but richness does not), likely due to growth of yeasts during early processing leading to altered species abundances compared to berries.

**Figure 4.**
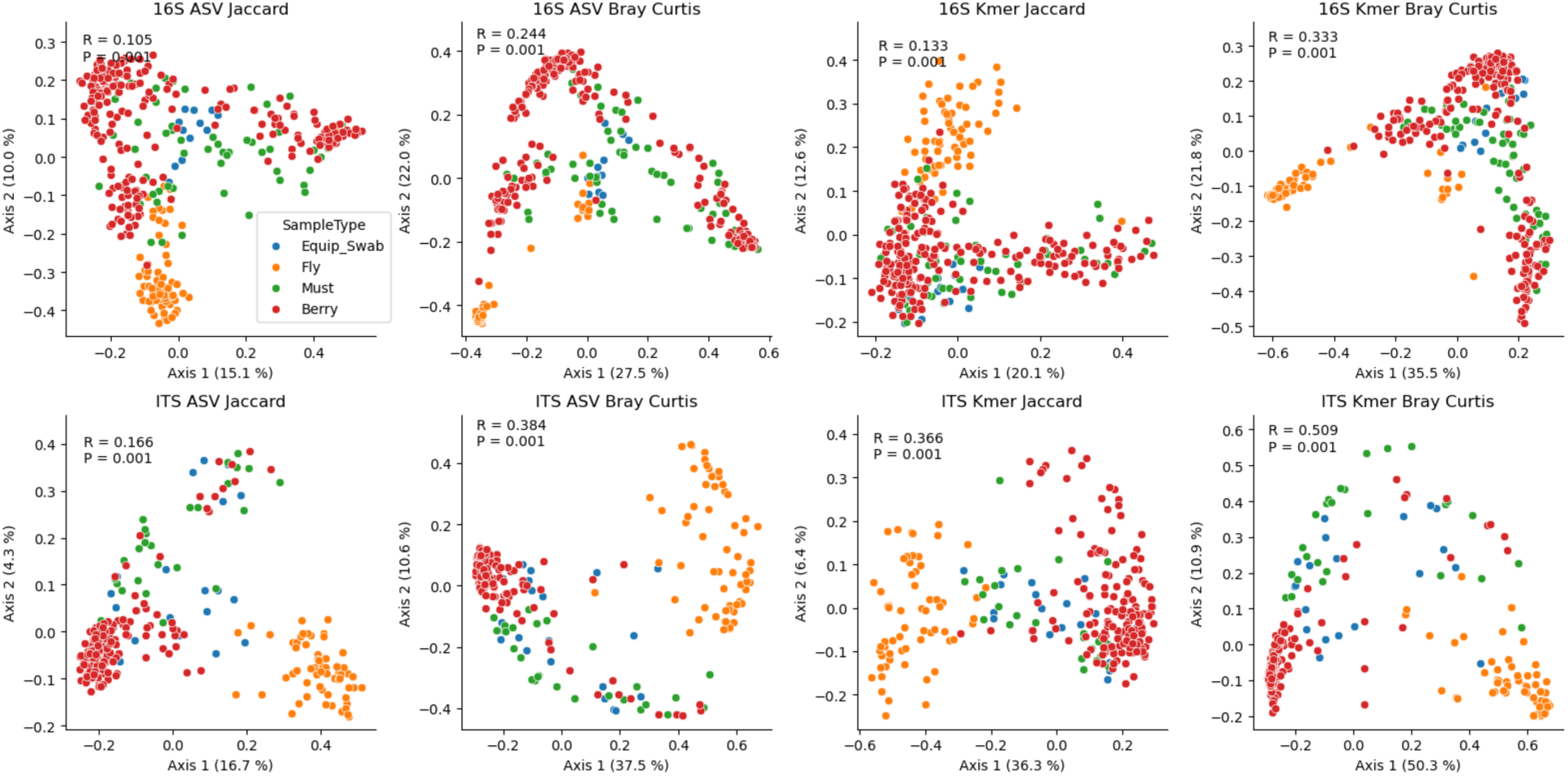
Vineyard and winery sample types exhibit significant differences in beta diversity. Jaccard distance PCoA Plots comparing bacterial (16S, top) and fungal (ITS, bottom) composition between grapes (red), grape must/juice (green), equipment swabs (blue), and fruit flies caught in the winery (orange). Inset: one-way PERMANOVA R2- and P-values.

**Figure 5.**
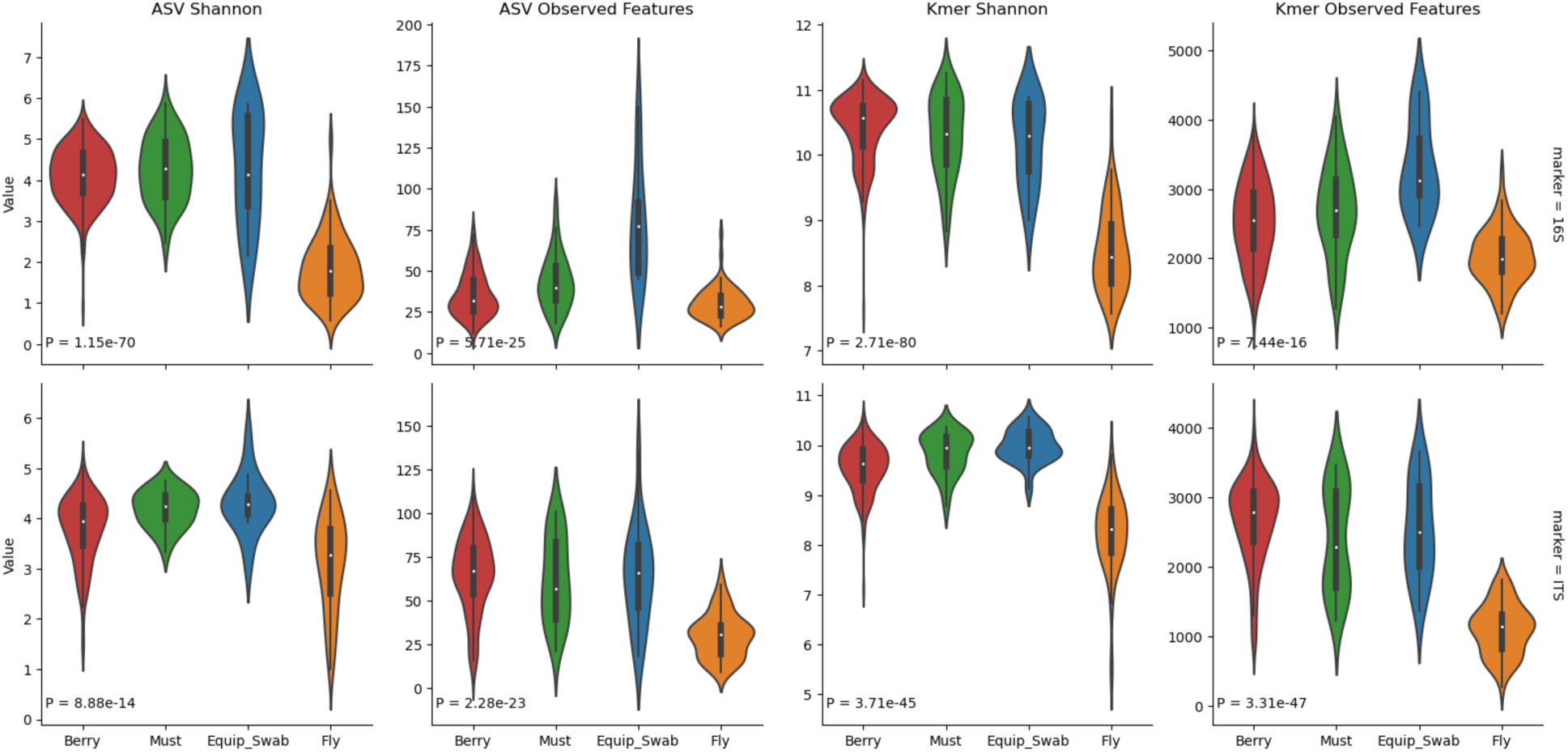
Alpha diversity varies significantly between vineyard and winery sample types. Shannon H and observed feature counts (richness) of ASVs (left two columns) and k-mers (right two columns) are shown for bacterial communities (top row) and fungal communities (bottom row) for grape berries (red), must/juice (green), equipment swabs (blue), and fruit flies caught in the winery (orange). P-values indicate the result of one-way ANOVA tests comparing all varieties.

Pairwise beta diversity distances (Figure 6) also show that the microbiota of musts are roughly equidistant to those of berries and equipment swabs, indicating that they exhibit intermediate compositions between the two sample types. This suggests that equipment contact causes assimilation between the microbiota of musts and equipment surfaces, and microbial transmission into musts during processing. However, almost all PERMANOVA p-values are significant (P < 0.05), indicating that within-group distances are lower than between-group differences; in other words, must samples are more similar to each other than they are to any other sample type, also suggesting an intermediate composition. Whereas berries have very low relative frequencies of fermentative yeast species including *Hanseniaspora* spp. and *Starmerella* spp., these are present at moderate abundances in both equipment and must samples (Figure S9). Must are also distinguished by higher mean relative abundances of the acetic acid bacteria *Gluconobacter* and *Komagataeibacter*, among others (Figure S9), suggesting that growth of aerobic bacteria already occurs during early fruit processing before fermentation is initiated.

**Figure 6.**
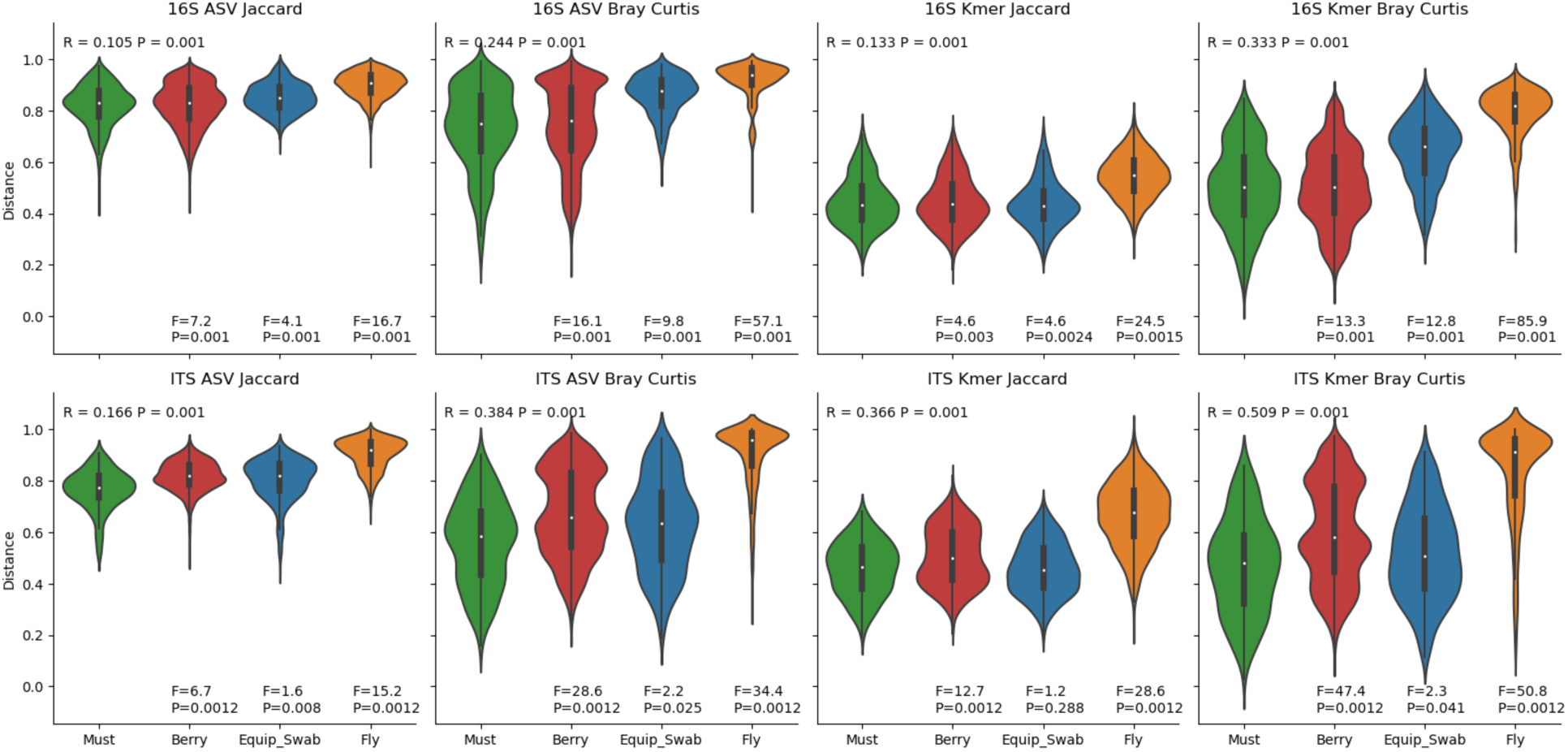
Beta diversity pairwise distances to must samples. Violin plots show the distribution of distances between must samples and each other sample type. PERMANOVA R-value and P-value for all group comparisons are displayed at the top of each panel; F-scores (effect size) and P-values for pairwise group comparisons are displayed at the bottom of each panel.

These findings are corroborated by network analysis of species observed at individual sites (Figure 7), showing that the species with higher weights (i.e. comprising more ASVs) shared exclusively between grapes and must correspond to plant-associated and environmental yeasts such as *Sarocladium zeae*, *Curvularia subpapendorfii*, and *Thelebolus globosus*, further indicating the relevance of vineyard taxa in these communities. Many other species are exclusively shared by must and equipment or flies, suggesting introduction within the winery. In contrast to the yeast-dominated grape-and-must-associated species, most of the nodes that are exclusive to must and equipment or flies are members of the Proteobacteria and Actinobacteria. Finally, many species are shared by all four sample types, which correspond to generalistic taxa. Overall the network shows a highly modular structure characteristic of biological networks.

**Figure 7.**
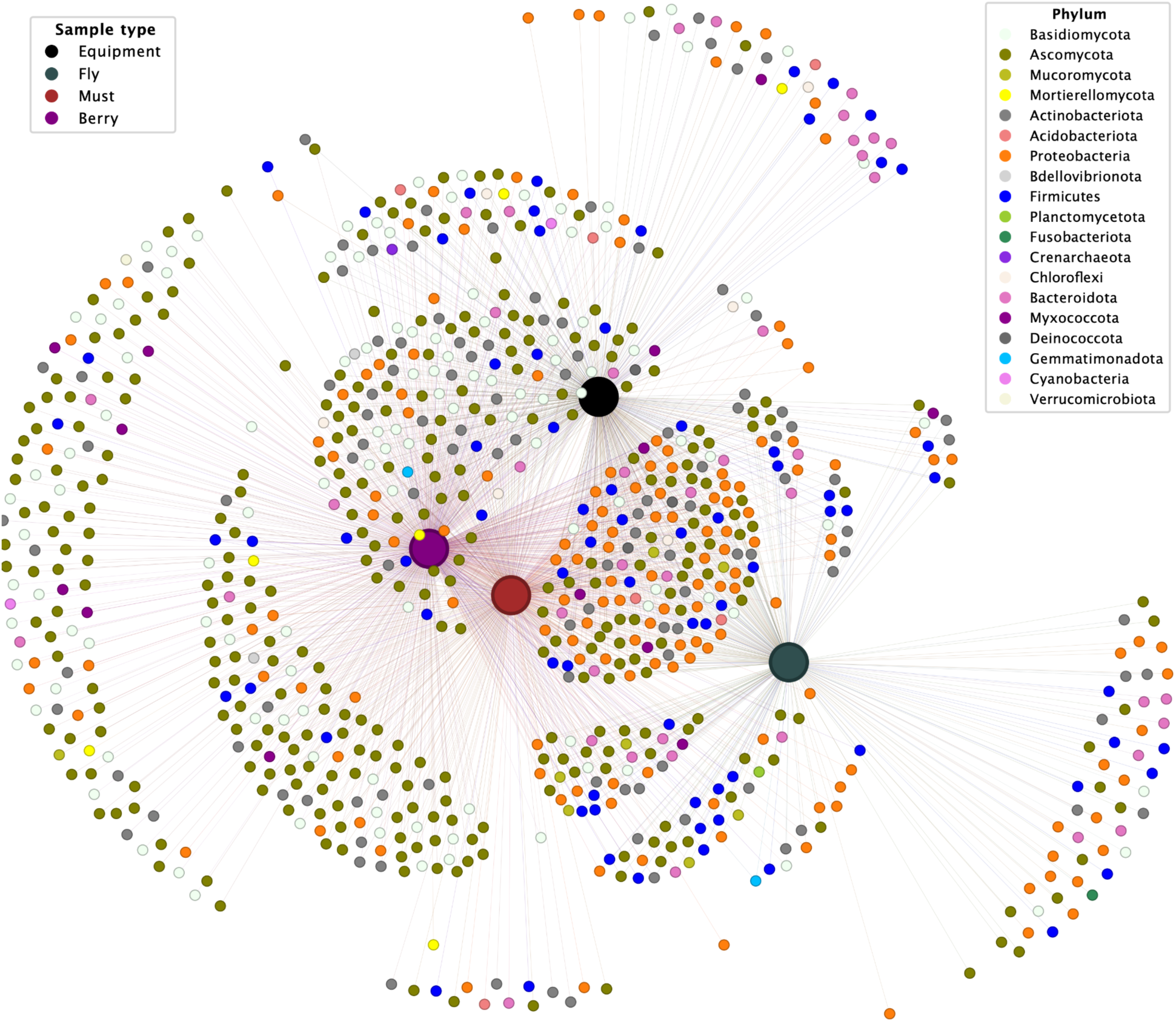
Bipartite network visualization of microbial species and sample types. ASVs were grouped by species and color-coded by phylum, while samples were categorized into four groups, represented by larger nodes: Berry (purple), Must (red), Fruit Flies (green), and Equipment Swabs (black). Smaller nodes represent species, with rescaled edges indicating their presence in each sample.

Random Forest classifiers trained to predict sample types demonstrate high predictive accuracy (accuracy = 0.897 ± 0.035; micro-AUC = 0.99; Figure S10), indicating large differences between all sample types. The top predictive features (Figure S11) indicate that predictive performance is driven by higher abundance of filamentous fungi in berries, musts and equipment swabs, and fermentative yeasts (including *Saccharomyces cerevisiae* and *Hanseniaspora uvarum* ASVs) associated with fruit flies and other sample types to a lesser extent.

### Fruit flies in the winery: a reservoir for fermentative yeasts?

The starkest contrasts in alpha and beta diversity were exhibited by fruit-fly-associated microbial communities (Figure 4-6). This was driven by particularly high relative abundances of fermentative yeasts associated with fruit flies (e.g., *Saccharomyces* spp. and *Hanseniaspora* spp.) and the obligate intracellular bacterium *Wolbachia*, which interestingly was also found at moderate frequency in both grape and must samples, reflecting insect interactions with the fruit (Figure S9). *Saccharomyces* spp. (Figure S12) were found almost exclusively in fruit flies, and many *Hanseniaspora* ASVs were predominantly detected in fruit flies, though other ASVs were frequently found in grapes, musts, and equipment, indicating that several different subpopulations of *Hanseniaspora* (represented by different sequence variants) were present in the vineyard and winery environments.

### Impact of processing on the grape must microbiota

The microbial communities associated with grapes evidently experience a subtle transformation during processing in the winery. As we collected grape must or juice samples at different stages of processing, this provided an opportunity to qualitatively examine how the microbial profile changed during processing. Samples from two separate lots of Chardonnay were taken from the crusher and at 10-minute intervals from the press, as the press cycle was programmed to increase in pressure in a stepwise manner every 10 minutes. This included free run juice prior to the press cycle initiation and juice samples from the settling tank immediately after pressing. The fungal profile progressively changed during this time course, exhibiting a gradual decline in relative abundance of the filamentous fungi *Alternaria, Erysiphe*, and *Stemphylium*, and a gradual increase in ASVs assigned to the yeasts *Hanseniaspora, Starmerella,* and *Lachancea* (Figure 8). No clear qualitative trends could be detected in the bacterial communities in the press samples (Figure S14). Replicate samples were not taken from these two lots, limiting statistical testing. Nevertheless, this demonstrates that the fungal community composition qualitatively changes during the course of pressing, and suggests that fermentative yeasts are able to begin growing during this time. This suggests that grape processing/pressing is an early window of opportunity during which time vineyard- and winery-resident microorganisms can initiate fermentation and potentially contribute to the chemical composition of the wine, providing an entry point for adventitious strains to influence wine quality even in wines that are later inoculated.

**Figure 8.**
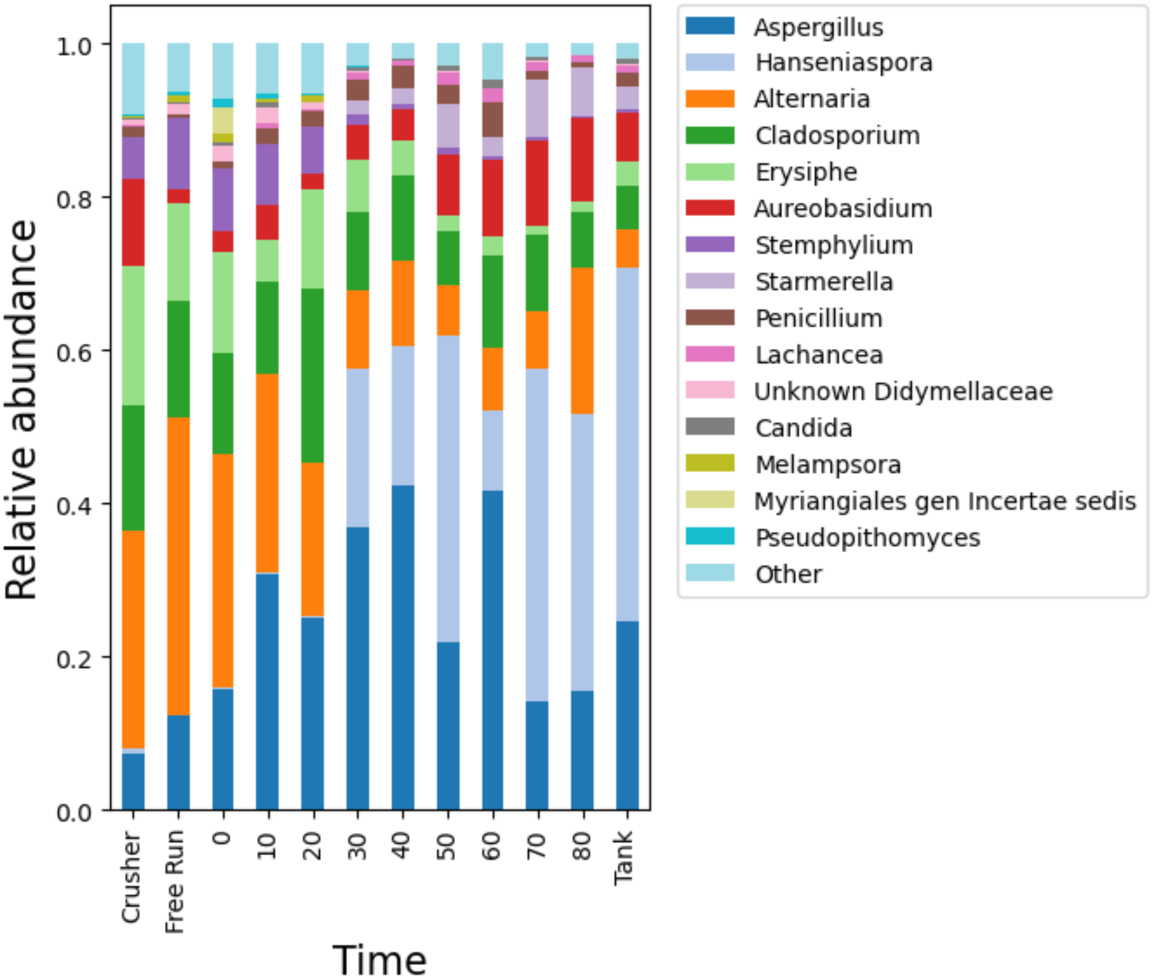
Dynamic changes in the fungal community profile of grape must samples during processing. Stacked barplots show the relative abundance of fungal genera in grape must samples at different press times, as well as crushed grapes and free run juice samples immediately prior to pressing, and settling tank samples immediately after. All taxa observed at < 0.01 relative abundance are binned into “Other”.

## Discussion

Parsing the drivers of microbial biogeography in wine production systems is inherently challenging. Thus far, the majority of studies investigating microbial ecology in vineyards have collected fruit from either single vineyards or from disparate vineyards, often managed differently, or planted to different varieties. The goal of the current study was to assess the effect of distance or site by comparing two vineyards planted to the same cultivar, farmed by the same group and in the same way, with no major geographic features separating the sites in an attempt to reduce covariates that drive regional differences between vineyards, especially at larger spatial scales. Embedded in this experiment was an attempt to track communities detected in the two vineyards through the pre-fermentation winemaking processes to understand their impact on community structure. Previous studies have shown that the winery is a potential source for adventitious microorganisms in wine fermentations [14], [15], [43], [44], but it is unclear how these organisms impact purported microbial terroir, or to what extent it changes the microbial community present.

The potential impact of cellar microbiota is likely different for different wineries based on cleaning practices, environmental features (e.g., materials and indoor conditions), and winemaking protocols. Hence, the results in this study are effectively a “case study” that may not generalize to all winemakers, but both vineyards in the current study were processed in the same winery on the same day to control for differences observed between wineries in previous studies. The microbiome data presented here demonstrate that crush equipment reflects contact with plant material during harvest, and that community structure changes across the fruit processing cycle from fruit samples to press fractions, showing that the winery environment can serve as a potential reservoir for microorganisms to transmit across batches and vintages to seed successive fermentations in a single winery.

Additionally, the present findings indicate that, although microorganisms from the winery environment clearly shed into grape must during processing, the microbial community profile from the vineyard remains quantitatively intact. Our findings show that two proximal Chardonnay vineyards retain their distinct microbial identities through processing, suggesting that vineyard site-specific microbiota patterns persist to the start of fermentation and can even grow during fruit processing prior to inoculation or sulfite addition, potentially contributing to the chemical/sensory characteristics of the resulting wine. The fruit and must samples each were differentiated by source vineyard when examining both fungal ITS and bacterial V4 data. The fruit samples from the two vineyards harbored distinct microbial communities, driven by unique ASVs, and differentially abundant taxa. This includes unique ASVs that were predominantly abundant in one or the other vineyard from wine-relevant genera, such as the many *Hanseniaspora* ASVs that were uniquely detected in Tyree but not in RMI. This is a startling difference, given that these are proximal vineyards (∼3 km apart), under uniform management, and sharing homogenous climate and soil conditions. This finding suggests that microclimate effects and other factors like land use context could play a role; the proximity of Tyree to a riparian zone is speculated as a possible factor, serving as a reservoir for transmission of microorganisms from the surrounding environment.

This study is also the first, to our knowledge, to show how the abundance of wine-relevant yeasts originating in the vineyard changes during fruit pressing. White wines in particular are subject to press fraction decisions due to the changing chemical composition of must as fruit is pressed. Here we show that pressing time also has a sizable impact on the microbial communities of grape juices as well. Unfortunately, only two batches were collected without replication of the press fractions, so this finding could not be statistically tested. Investigating how separating press fractions or adjusting press times alters the microbiological and chemical quality of wines remains an interesting question for follow-up.

Previous studies have shown that grape varieties harbor distinct microbial communities within and between regions [1], [2], but it has rarely been demonstrated at the intravineyard scale [45], [46]. The differences previously reported are thought to be the result of morphological differences in the clusters, skins, and ripening curves of different varieties resulting in distinct surface conditions affecting microbial life. However, as each variety was planted to a different block in the Tyree vineyard it remains a possibility that the block-level differences observed between varieties are influenced by spatial effects, e.g., dispersal limitation effects within the vineyard. Further work should attempt to parse these complications with appropriate study design, e.g., random complete block design, which is challenging given the time-to-maturity for grapevines. In the current study, the observation that varieties cluster by shared parentage (Cabernet Sauvignon with Colombard, Grenache Blanc with Grenache Noir), strengthens the likelihood that genotype effects are at play, rather than block and dispersion effects alone, but additional work is required to parse these effects.

Another interesting observation in this work was the apparent exchange of microorganisms — especially fermentative yeasts — between fruit flies captured in the winery environment, grapes, and grape musts. Fruit flies have an intimately intertwined ecological relationship with yeasts in the natural world, and in winery and vineyard ecosystems fruit flies may be important vectors for yeast transmission to spontaneous fermentations and to fruit surfaces pre-harvest [47]. Yeasts are commonly vectored by insects between seasonally ephemeral sugar sources, an ecological relationship that has been described as the ‘dispersal-encounter hypothesis’ [48]. Facilitating this interaction in fruit flies is the preferential attraction of fruit flies to yeast-produced volatiles such as ethyl acetate and isoamyl acetate over fruit metabolites [49]. Quan and Eisen [47] previously characterized the fungal microbiota of fruit flies in a winery environment, demonstrating spatial differentiation and carriage of diverse yeasts, an observation that we corroborate in our study. Furthermore, we characterized the bacterial microbiota of winery-associated fruit flies (which was not described by Quan and Eisen); our findings suggest that fruit flies can also be vectors of bacterial contaminants within winery environments, and the presence of the obligate intracellular bacterium *Wolbachia* in grape and fruit samples serves as a marker of fruit fly contact. Moreover, the high frequency of filamentous fungi, e.g., *Fusarium*, *Aureobasidium*, *Alternaria*, *Erysiphe*, and *Aspergillus*, suggest that fruit flies may be vectors for other fungi beyond filamentous yeasts. Furthermore, a large percentage of both fungal and bacterial ASVs were shared between winery equipment and fly samples, suggesting either co-localization [47] or interaction.

Some limitations of this study are worth noting, and inform points of improvement for future studies. First, the commonly used 16S rRNA gene primers employed here also amplify chloroplast and mitochondrial 16S rRNA genes, leading to low bacterial sequencing efficiency 16S rRNA gene dataset; in the future, use of blocking primers, PNA clamps, or similar approaches could be used to improve efficiency [50], [51]. Second, even though we selected two proximal vineyards growing the same grape variety (Chardonnay) under uniform management practices, some partial differences were still apparent between these vineyards, and hence are partial covariates. These include: (i) varying rootstocks, which are known to impact the rhizosphere microbiota [52], [53]; (ii) only partial overlap in soil series composition (as Tyree contained a mixture of soil types), which could also be expected to alter the soil microbiota and although impacts on the grape microbiota are unclear, the soil serves as an important reservoir for microorganisms that colonize the grape surface [13], [16]; (iii) partially different use of grapevine clones, which could be more likely to directly impact the grape microbiota on the fruit surface; and slight differences in vineyard row spacing, the impact of which is unknown but could theoretically alter the grape microbiota via microclimate effects. These may all partially contribute to the strong site effects observed here, though all of these are not expected to have such a significant impact on the grape microbiota on their own. Nevertheless, these factors deserve further consideration in future work that could isolate and directly examine the impacts of these and other site-specific factors on microbial community colonization of grapevines. Finally, grapes from different varieties were collected just prior to harvest so that all were collected at similar degrees of ripeness; this means that different varieties were collected on different sampling dates. Given the uniformly warm, dry weather conditions present in central California during August and September, we do not anticipate that the microbiota would change significantly during this time period and expect that degree of ripeness (sugar concentration) is a more important factor to control.

## Conclusion

Here, grapes from two nearby vineyards were sampled to examine site-specific effects on the grape-associated microbiome, and how this microbiome changes during fruit processing. Chardonnay grapes were sampled on the same day from each vineyard and found to harbor distinct fungal and bacterial communities in spite of the close proximity, similar climatic and edaphic conditions, and uniform farming practices of these two vineyards. Controlling for spatial distance, management type, variety, and clone in this study furthers our understanding of the factors that shape microbial assembly in vineyards and wine fermentation, demonstrating that these factors are significant but alone do not explain the vineyard site-specific microbial community signatures observed in previous studies [1], [2], [3], [8].

Furthermore, these differences persisted through processing in a single winery. Microbial community structure was shown to change across fruit processing and was assessed for the first time in conjunction with crush equipment in a working winery. This study does not explicitly address the origins of microbes on fruit, or why these vineyards located at a close distance, planted to Chardonnay, harbor distinct communities, but rule out the possibility that subtle differences in vineyard management explain these differences. We show that microbial diversity and community structure between vineyards and across sample types differed for fungal and bacterial data, congruous with previous findings, yet at a smaller scale. While comparatively fewer fungal than bacterial taxa were present, communities separated more clearly in ordinations, and distinguishing taxa between vineyards were distinct at the sub-genus level. Bacterial diversity on the other hand was reflected in phylum-level distinctions in relative abundance and each vineyard harbored unique taxa. Given the similar mutability of soil bacterial communities at small spatial scales [54], [55], this re-broaches the possibility that soil is a prevalent reservoir for bacterial epiphytes in grapevines and other plants [7], [56], [57] and spatial variations in soil microbiota composition could explain some spatial variations in phyllosphere communities. Both findings represent the interplay between similar ecological niches and dispersal limitation that make up the previously reported phenomena of microbial terroir.

The presented evidence supports the conclusion that fruit is the dominant source of microorganisms in grape musts under normal cleaning conditions in a production winery, though winery equipment and fruit flies can also be significant reservoirs, particularly for fermentative yeasts. Vineyards planted in close proximity to one another can harbor distinct microbial communities, and these communities or what is detected changes during processing, both at the crusher, and across press fractions. The crush equipment is likely a two-way medium for microbial transfer and reflects contact with plant material as previously reported, and appears to contribute to grape must microbiota. Even so, microbial communities, distinct by vineyard, retain their identity from the fruit samples to must, underlining the strong potential of vineyard-specific microbial communities to contribute to the quality characteristics of regional wines.

## Acknowledgments

NAB gratefully acknowledges financial support from the Swiss National Science Foundation [310030_204275]. RGG was funded by the Wine Spectator Scholarship, Louis R. Gomberg Scholarship, Adolf L. & Richie C. Heck Research Fellowship, Mario P. Tribuno Memorial Research Fellowship, Jastro Shields Research Award, Michael Bonaccorsi Scholarship, Knights of the Vine Scholarship, Mangahas Family Scholarship, and the Paul Monk Scholarship whose generous support of UC Davis made his work possible. The authors thank Morgan Bennett (UC Davis) for assistance with sample collection and DNA extraction. The authors also thank the many vineyard and winery workers in the Department of Viticulture and Enology at UC Davis who indirectly contributed to this work through routine management of the vineyards and winery studied herein.

## Author contributions

NAB and DAM conceived and designed the study. MS and NAB collected samples. RGG, NAB, LF, and RHV analyzed the data. NAB and RGG wrote the manuscript, with contributions from other authors. DAM and NAB acquired funding for this work.

